# Spatial and directional tuning of serial dependence for tracking eye movements

**DOI:** 10.1101/2024.06.03.597076

**Authors:** Alexander Goettker, Emma E.M. Stewart

## Abstract

An attractive influence of past sensory experience on current behaviour has been observed in many domains, such as for perceptual decisions and motor responses. However, it is unclear what sort of information is integrated across trials, and the limits of this integration, especially for oculomotor behavior. Here we provide a detailed and systematic investigation of the spatial and directional tuning of serial dependence for oculomotor tracking. In a series of experiments, we measured oculomotor responses to sequences of movements: the first movement (the prior) could move at different velocities (5 or 15 deg/s), and could additionally vary in its spatial location or direction relative to the following movement. The second movement (the probe) always moved at the same velocity (10 deg/s) and was constant across all experiments. We observed that eye velocity for the probe movement was faster when following the fast prior compared to following the slow prior, replicating attractive serial dependence. Importantly, this effect stayed consistent for distances of up to 30 deg between probe and prior, strongly suggesting a retinotopic coordinate frame. When we manipulated the direction of the prior, we observed that the strength of the serial dependence on eye velocity as well as eye direction was modulated by the relative angle between prior and probe. We observed stronger serial dependence for prior directions more similar to the probe direction. The strength of the effect on eye velocity and eye direction was correlated, suggesting a shared mechanism controlling these effects. Across all experiments, we observed that even when the prior moved in the opposite direction to the probe, there was a residual attractive effect. This suggests that serial dependence for oculomotor tracking consists of two components, one retinotopic, direction-tuned component and one more general component that is not direction-specific.

## Introduction

The perceptual and oculomotor systems use statistical regularities in the world to predict what will occur next, typically leading to an attractive serial dependence effect (Fischer & Whitney, 2014). The influence of past information on perception has been observed across a wide range of tasks, from ones related only to simple features to tasks with more complex, naturalistic stimuli (Cicchini et al., 2024; Manassi et al., 2023; Pascucci et al., 2023). Such serial dependency effects are not only observed in the perceptual realm, but also for different types of motor behaviors, ranging from eye (Cont & Zimmermann, 2021; Darlington et al., 2018; Deravet et al., 2018) to head movements (Zimmermann, 2021). A critical outstanding question across all serial dependence research is what kind of information gets integrated with current sensory input. While this question has been heavily studied for perceptual tasks (Manassi et al., 2023), the kind of information that is integrated, and the limits of this integration are still not well characterized in the motor realm.

For perceptual serial dependence effects, a recent meta-analysis (Manassi et al., 2023) demonstrated that for serial dependence to occur, past information needs to be 1) presented at a similar spatial location; 2) temporally close; 3) similar in a relevant feature; and 4) this relevant feature should be attended. Focusing on the spatial location, the integration of previous experience for perception seems to operate over a large spatial distance of up to 20 degrees (Manassi et al., 2023), forming a “continuity field” inside which information gets integrated (Fischer & Whitney, 2014). However, there is divergent evidence as to how spatial proximity is defined for perceptual serial dependence. Does the previously seen information need to be at a similar retinal location (retinotopic coding) to the current sensory information, or in a similar spatial location (spatiotopic coding)? Studies trying to dissociate spatiotopic and retinotopic effects in serial dependency find mixed evidence: While one study suggests that serial dependence operates in a large, retinotopic window (Collins, 2019), other evidence points to a spatiotopic mechanism (Mikellidou et al., 2021).

For oculomotor control, previous work has often focused on how previously seen target motion in one trial influences eye velocity in the following trial, and has largely ignored the type of information that is integrated. These studies have observed similar attractive serial dependence effects to perception: Faster motion on a previous trial (prior) led to a faster oculomotor response in the following trial (probe). These studies primarily manipulated the reliability of either the prior or the probe stimulus (Darlington et al., 2017, 2018; Deravet et al., 2018; Goettker, 2021), and showed that sequential effects were stronger when either the information in the previous trial was more reliable, or when the following sensory information was less reliable - consistent with reliability-weighted integration (Ernst & Bülthoff (2004); see Pascucci et al. (2023) for a discussion of similar perceptual results in serial dependence). Unlike for perception however, the characteristics of the stimuli that are varied for oculomotor control tasks are relatively limited, which makes it unclear what sort of stimulus information is critical for the integration of previous information. Darlington and colleagues (2017) replicated the temporal tuning evident in perceptual serial dependence (reviewed in Manassi et al., 2023) for oculomotor behavior, with lower serial dependence effects for stimuli that were further apart in time. They also manipulated target direction in addition to target speed: using different target directions, they showed that the initial eye direction in the following trials showed an attractive serial dependence effect. A recent study by Goettker and Stewart (2022) showed that the relevant feature space for stimulus integration is unrelated to object identity. They observed similar serial dependence effects independent of whether the prior and probe stimulus were the same (two Gaussian Blobs) or different (a depiction of a car and a Gaussian Blob). In addition, and in contrast to perception (Cicchini et al., 2021), they observed that serial dependence for oculomotor control is mediated by low-level retinal signals, ignoring contextual information. This also provides evidence for differences in the processing of sensory information for oculomotor control and perception (Spering & Carrasco, 2015), and highlights the need for more detailed studies of serial dependence for oculomotor control, since results cannot be directly transferred or generalized from the purely perceptual realm. While serial dependence for perception seems to match that for oculomotor control for some stimulus properties (i.e. temporal tuning, Darlington et al. (2017)), this may not be the case for all stimulus properties (i.e. identity, Goettker and Stewart (2022)).

The goal of this study was to provide a more detailed investigation of spatial and directional specificity for serial dependence for tracking eye movements. In the first set of experiments, we wanted to investigate whether the integration of information occurs in retinotopic or spatiotopic coordinates. We systematically varied the spatial locations of the stimuli while measuring sequential effects for target speed and target direction. In the second set of experiments, we wanted to provide a detailed investigation of the directional tuning of serial dependence effects. By presenting the prior stimulus moving in multiple different angular directions, we were able to compute how the strength of serial dependence is modulated by the angular difference between prior and probe.

## The spatial tuning of serial dependence for oculomotor behavior

To test the spatial tuning of the serial dependence effect for oculomotor tracking behavior, we designed a total of three different experiments. In each experiment, each trial consisted of two target movements: first the prior and then the probe (see for example Figure 1). The probe always moved with the same velocity (10 deg/s), however across the experiments the prior could differ in speed as well as its location and direction relative to the probe. Due to the matched sensory information during the probe trial, any systematic difference observed can be attributed to the influence of the prior. Given that the foveated tracking of a moving target ensures that the retinal target motion is the same across all conditions, we wanted to dissociate retinal from spatiotopic influences in serial dependence. We therefore compared conditions where the probe was either at the same or different spatiotopic location to the prior. In experiment 1, observers fixated on a small fixation dot either at the center or in the bottom half of the screen (8 deg vertical distance). The prior movement was presented at one of these two positions. To assess the influence of serial dependence, we measured the oculomotor response during the probe trial, in which the stimulus always moved at the same speed (10 deg/s) and was always presented at the central location, and was therefore either at the same or at a different spatiotopic location to the prior (see Figure 1A). Given that serial dependence for perception only integrates information up to 20 deg of distance between probe and prior (Manassi et al., 2023), we added the control experiments 2 and 3. Here a different group of observers were tested with distances of either 15 deg or 30 deg between the prior and probe (see Figure 1A). Due to limitations in vertical screen size, we changed the target movements to be vertical and introduced a horizontal position shift. To investigate the spatial tuning, we looked at how the spatial location of the prior affected serial dependence. To quantify this, we calculated the difference in eye velocity in the probe trial between fast and slow prior movement conditions (see Methods for more details).

**Figure 1.**
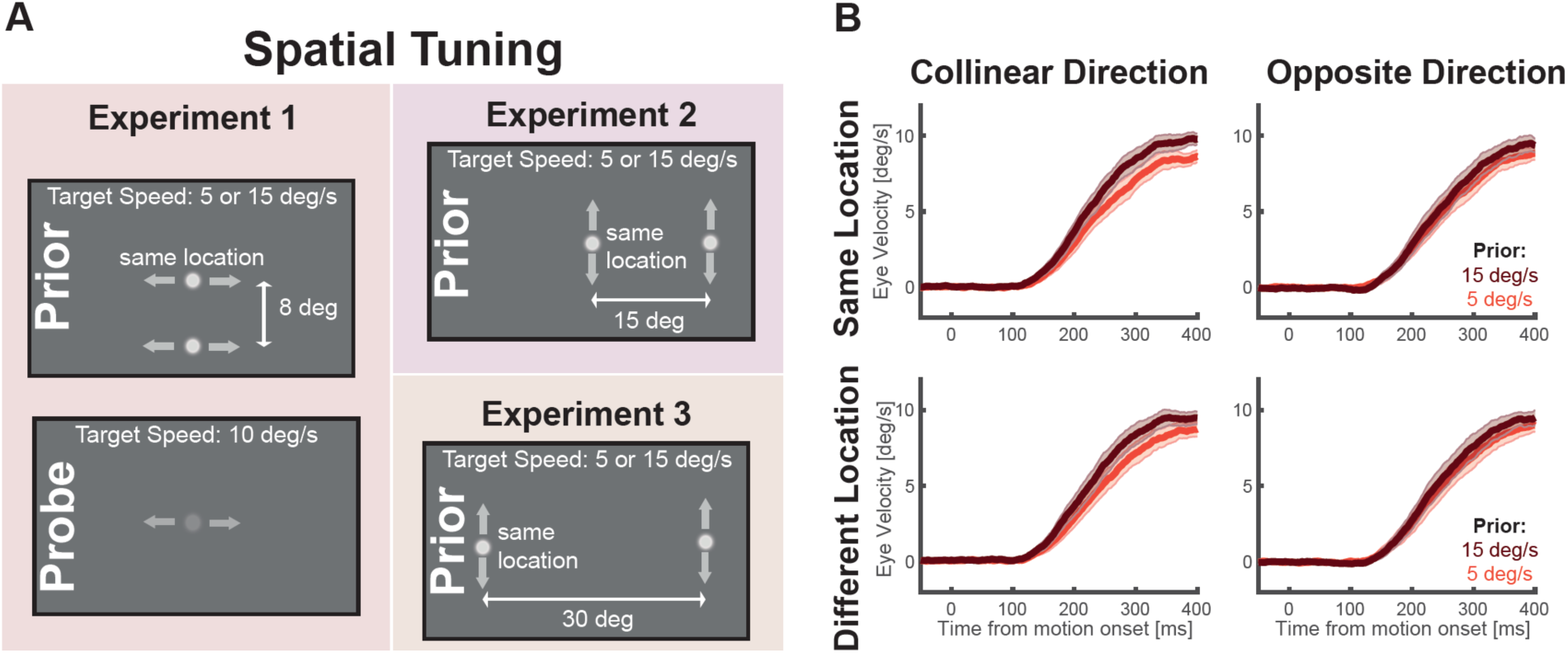
Overview of experiments for spatial tuning. **A** Depiction of the paradigms used for the spatial tuning experiments. Please note that while for experiments 2 & 3 only the prior movement is depicted, in all experiments a prior movement was followed by a probe movement. The left shows the conditions for experiment 1, where the prior and probe moved horizontally either to the left or to the right of the initial location. One prior movement was always identical to the location of the presented probe movement, while another one was presented at a different spatial position. For the prior movement, the target could move at either 5 or 15 deg/s, whereas the probe movement always moved at 10 deg/s. **B** The average eye velocity for probe trials for the four conditions presented in experiment 1. The different panels depict the different combinations of a collinear or opposite direction of the prior with respect to the probe and the same or a different starting location. The different shades of red indicate whether the prior moved with 5 or 15 deg/s. The shaded areas represent the 95% CI across observers.

### Methods - Spatial tuning

#### Observer characteristics

A group of 18 observers (mean age: 24.94 years; range: 19-56; 13 female) took part in experiment 1. A second group of 15 observers (mean age: 29.00; range: 25-38; 12 female) took part in experiments 2 & 3. Sample size was similar to previous work on serial dependence for oculomotor behavior (Deravet et al., 2018; Goettker, 2021; Goettker & Stewart, 2022). All participants reported normal or corrected-to-normal vision and were naïve with respect to the study. Experimental procedures were in line with the declaration of Helsinki and were approved by the local ethics committee. Written informed consent was obtained from each participant.

#### Setup

Participants sat at a table in a dimly illuminated room with their head positioned in a chin and forehead rest. Their eyes were roughly aligned with the upper half of a monitor (60 cm x 32 cm, 3840 x 2160 pixel, Phillips, Amsterdam, Netherlands) at a distance of 70 cm. Under these conditions the monitor spanned approximately 49 x 26 deg of visual angle. The experiment was programmed and controlled with Matlab 2020a (MathWorks, Natick MA) using Psychtoolbox (Kleiner, Brainard & Pelli, 2007). Gaze was recorded from one eye with a desk-mounted eye tracker (EyeLink 1000 Plus, SR Research, Kanata, ON, Canada) at a sampling frequency of 1000 Hz. To ensure accurate recordings before each block, a nine-point calibration was performed, and additional drift corrections were used at the start of each trial.

#### Experimental conditions

The goal of the first set of experiment was to assess the reference frame of the serial dependence effect for tracking eye movements. A single trial always consisted of two consecutive movements that observers needed to track with their eyes: the prior and the probe.

In *Experiment 1*, the prior movement differed across three dimensions: 1) it could start either in the center of the screen or vertically shifted downwards by 8 deg; 2) it could move to the left or to the right; and 3) could move either at 5 deg/s or at 15 deg/s. The probe movement always started in the center of the screen and moved at 10 deg/s, but could move to either the left or right. By comparing the influence of the prior on the probe trials across these different manipulations, we could assess whether the influence on the probe trial differed depending on the different starting location of the prior. Since participants always were tracking the movement with their eyes, the retinotopic movements were comparable across all conditions, whereas due to the different starting locations, the movements differed in their spatial coordinates.

Each trial started with a drift correction, where participants could start the trial by looking at the fixation cross and pressing the space bar in a self-paced manner. Then, a small red fixation dot (diameter = 0.2 deg) was presented for a random time between 1 and 1.5 s. The small fixation dot appeared at the starting location of the prior movement (either in the center or shifted downwards by 8 deg). After the random fixation time, the fixation point disappeared and the target stimulus, a white Gaussian blob (SD = 0.4 deg, Contrast = 1) appeared and moved either left or right, at either 5 or 15 deg/s. The target always appeared in a step-ramp manner, so it crossed the initial fixation dot after 200 ms (e.g., when the target moved to the left with a speed of 15 deg/s, it appeared 3 deg to the right of the fixation dot). The target moved for 1 s and then the fixation point for the probe movement immediately appeared. After a new random fixation time between 1 and 1.5 s, the probe movement was presented in the same fashion as the prior movement. To enhance the serial dependence effect, the probe stimulus was a lower contrast Gaussian blob (SD = 0.4 deg, Contrast = 0.1). Overall observers completed 128 trials (2 starting locations * 2 prior directions * 2 prior target speeds * 2 probe directions * 8 repetitions) in one block, and completed a total of 3 blocks. Each block lasted between 15-20 minutes.

In *Experiments 2 and 3*, the timing of and order the stimulus presentation were identical: the prior could either move at 5 or 15 deg/s and the probe always moved at 10 deg/s. The major difference now was that the location and direction of the target movements shifted. To allow for larger distances between the prior and probe stimulus within the limits of the screen movements became vertical instead of horizontal. In experiment 2 the prior stimulus either moved upwards or downwards and could do so either starting from the center of the screen or shifted 15 deg to the left. In experiment 3, the prior stimulus also either moved upwards or downwards and could start at either shifted 15 deg to the left or 15 deg to the right. In both experiments, the probe always moved either upwards or downwards and always was presented at 15 deg to the left. In this way in Experiment 2 there was a condition with 15 deg distance between the prior and the probe and in Experiment 3 a condition with 30 deg distance between prior and probe. Observers again completed 128 trials (2 starting location * 2 prior directions * 2 prior target speeds * 2 probe directions * 8 repetitions) per block and two blocks per experiment. Each block lasted between 15-20 minutes.

##### Exclusion criteria

Trials were excluded from the analysis if (1) there was more than 500 ms missing data during the prior or the probe movement, (2) if during the critical interval of the probe trials (−50 to 400 ms after motion onset), there were residual velocities larger than 30 deg/s (three times the target speed), indicating a potentially missed corrective saccade, (3) if in the probe trial no valid pursuit onset was detected or the latency was below 50 ms or larger than 400 ms. Based on these criteria we could include 6764 out of 6912 trials (98%) for experiment 1. Since vertical pursuit is less accurate, for experiments 2 and 3, trial exclusions were slightly higher. For experiment 2 we had 3 observers with more than 30% invalid trials, which were excluded from the statistical analysis. For the remaining observers we included 2731 out of 3072 trials (89%) for the analysis. For experiment 3, we again had to exclude 3 observers due to more than 30% invalid trials and included 2697 out of 3072 trials (88%) for analysis.

##### Analysis

Eye movement data were digitized on-line and analyzed off-line using Matlab software. First, blinks were linearly interpolated (from 30 ms before detected blink onset to 30 ms after detected blink offset) and the eye position was filtered with a second-order Butterworth filter, with a cutoff frequency of 30 Hz. Then eye velocity was calculated as the first derivative of the filtered position traces. Saccades were identified based on the EyeLink criteria with a speed and acceleration threshold of 30 deg/s and 4000 deg/s^2^, respectively. After the detection of saccades, a linear interpolation of the eye movement velocity around the time of the saccade (from 30 ms before detected saccade onset to 30 ms after detected saccade offset) was performed. Eye movement velocity was filtered with an additional low-pass Butterworth filter with a cutoff frequency of 20 Hz. We also detected pursuit onset in the probe trials as the first point where the horizontal eye velocity reached 30% of the target speed and remained at this speed for at least 100 ms.

To compute the influence of the prior, all velocity traces during the probe were aligned to target movement onset, and if the probe was moving to the left, the horizontal eye position was multiplied by -1 to align it with the rightwards movements. Then the average *horizontal* velocity (for experiment 1) or *vertical velocity* (for experiments 2 and 3) in the probe trial was computed across all different prior conditions (different prior target location, relation between prior and probe direction). The strength of the serial dependence effect was then quantified by computing the difference between the eye velocity profiles between the fast and slow prior movements. To do this we used the summed difference in the analysis window of 100 to 400 ms after motion onset normalized by the number of frames. In this way, we obtained a measurement of serial dependence for each of the different prior conditions and each observer.

To estimate the contribution of the unspecific component of the serial dependence effect, for each experiment, we computed the mean of the conditions where the target moved in the same or in the opposite direction. Then we divided the mean of the effect for the opposite direction by the mean of the effect in the collinear direction. To see how stable the effect of the unspecific component is, we used bootstrapping to estimate the variance of our estimate. To estimate the variance, we took the data from all valid subjects and took 20 random samples with replacement 1000 times, and repeated the procedure mentioned above.

### Statistical Analyses

T-tests were conducted in Matlab 2021b. Linear mixed models were computed using the R package nlme (Pinheiro et al., 2023; Pinheiro & Bates, 2000).

## Results - Spatial tuning

Eye velocity in the probe trial was affected by the speed of the prior (slow, 5 deg/s, or fast 15 deg/s), the starting location of the prior (same or different), and the relative direction of the prior (collinear or opposite as the probe; Figure 1B shows the raw eye velocity for Experiment 1). To quantify and visualize the serial dependence effect, for each experiment, we computed the difference in eye velocity for the probe between the fast and slow prior and investigated how this was modulated by the starting location and relative direction of the prior (Figure 2). In this way, positive values indicate an attractive effect (faster prior è faster eye velocity in the probe), whereas negative values indicate a repulsive effect (faster prior è slower eye velocity in the probe).

**Figure 2.**
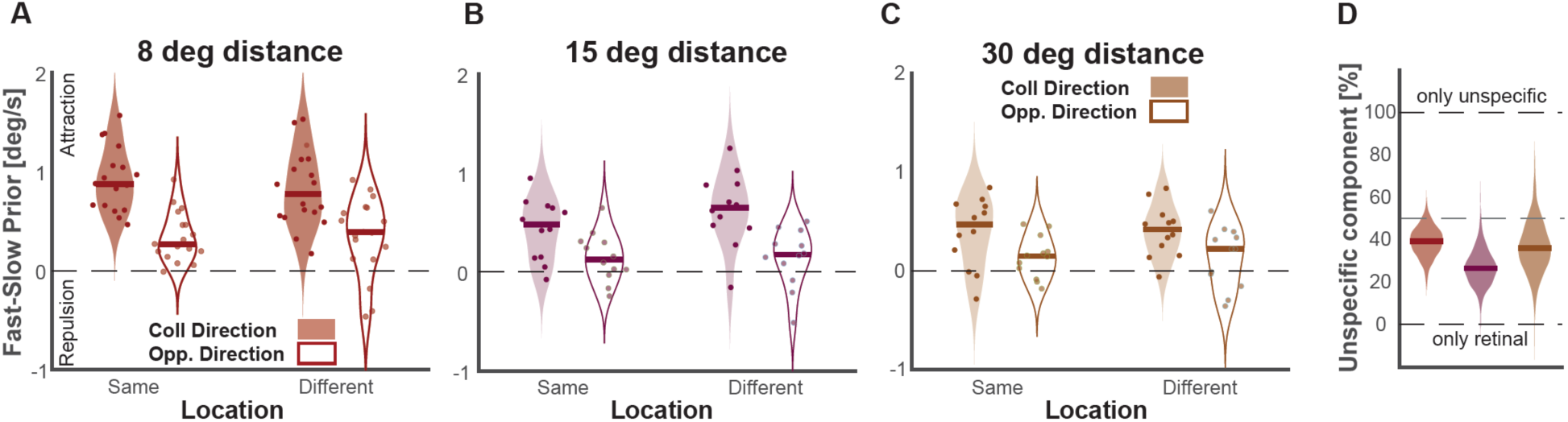
The spatial tuning of serial dependence. The strength of the serial dependence effect quantified as the difference in eye velocity in the probe movement depending on the prior target speed. **A** shows the results of experiment 1, **B** shows the results of experiment 2, **C** shows the results of experiment 3. Please note that these results are not directly comparable since they were obtained with a different group of observers and horizontal target movements for experiment 1 and vertical target movements for experiment 2 & 3. The figure shows violin plots depicting the distribution of the effect, the solid lines depict the median. Dots depict the individual observers. Positive values indicate an attractive serial dependence effect, negative values a repulsion effect. The plots are split depending on the prior target location (left vs right on the x-axis) and the relative direction of the target movements (collinear: filled distributions, opposite direction, open distributions). **D** The relative contribution of the direction unspecific component. Distribution estimated via bootstrap. See text and methods for more details.

To test the effect of prior location and direction across the three experiments (1,2, and 3), we used a linear mixed model. We specifically wanted to test how the serial dependence effect for each participant depended on distance between prior and probe (0, 8, 15, 30), collinearity between the prior and probe (collinear or opposite), and eye movement direction (horizonal or vertical). We included a random effect of participant (with random intercepts and slopes for collinearity conditions). The full model specification was: serial_dependence_effect ∼ prior_colinearity * prior_distance * eye direction + (∼1|subject/prior_colinearity). The magnitude of serial dependence was influenced by the collinearity between the probe and prior: F(1,30) = 61.27, p <0.0001; and by whether the movement direction was horizontal or vertical: F(1,30) = 21.82, p < 0.001; however it was not influenced by the distance between probe and prior: F(1,100) = 0.98, p = 0.32, nor were there any interactions between collinearity, distance, and eye direction. Our results provide evidence for a retinotopic reference frame for oculomotor tracking: the strength of the serial dependence effect was larger if prior and probe moved in the same direction, but this effect was independent of the distance between prior and probe. The main effect of movement direction is not surprising, given the well-documented differences in overall tracking performance, with more accurate tracking for horizontal movements (Ke et al., 2013). Crucially, there was no interaction between movement direction and the other factors, indicating that the retinotopic pattern of results generalized across experiments.

To our surprise, there was still a significant serial dependence effect even when the probe and prior moved in opposite directions. A one-tailed t-test against 0 (Bonferroni corrected for three multiple comparisons, p=0.05/3 = 0.167) revealed that for Experiment 1 (t(17) = 5.44, p < .0001), Experiment 2 (t(12) = 2.54, p = .014), and Experiment 3 (t(11) = 2.623, p = .012), there was a significant attractive effect for conditions where the probe moved in the opposite direction to the prior (averaged across the different spatial locations). The effect was highly systematic, as most participants still showed an attractive effect even when the prior moved in the opposite direction (see positive values for the open distributions in Figure 2). This suggests that the serial dependence effect we measure contains two different components: One component that is direction-specific and seems to be retinotopic, and another component that seems to be a more general component that is neither spatially-nor direction-selective. To quantify the importance of this direction-unspecific component we computed the mean serial dependence effect for the targets moving in the opposite direction and normalized it by the mean effect for targets moving in the same direction (see Methods for more details). Across all experiments, the magnitude of the direction unselective effect was between 25% and 40% of the effect measured when the targets moved in the same direction (39% for experiment 1; 26% for experiment 2; 35% for experiment 3).

In summary, for spatial tuning, we observed clear evidence for a retinotopic coding, as the serial dependence effects were consistent regardless of the spatial location the prior. The strength of the effects stayed consistent up to a distance of 30 deg, which is significantly larger than the spatial tuning described for perceptual effects (Manassi et al., 2023), and emphasizes the importance of retinal motion as the main factor underpinning serial dependence for oculomotor control (Goettker & Stewart, 2022). In addition to a direction-tuned retinal component, we observed a second component, which resulted in significant attractive effects even when the prior and probe moved in opposite directions, and irrespective of the distance between them. While the overall effect size differed between observers and experimental contributions, the relative contributions of the retinal and direction unspecific components stayed comparable across the different experiments and target distances.

## The directional tuning of serial dependence for oculomotor behavior

In the spatial tuning experiments, we observed a clear retinal tuning that was modulated by the movement direction of the prior relative to the probe. In the second series of experiments, we were interested in investigating this feature tuning in more detail (see Figure 3A). We designed two further experiments, where we systematically varied the movement direction of the prior relative to the probe. In these experiments, the probe movement always started in the center and moved at 10 deg/s to the right, whereas the prior started at the same location but could move in one of eight different directions and two speeds: either slow (5 deg/s) or fast (15 deg/s). We shifted the distribution of prior directions between experiment 4 (−30°, -15°, 0°, 15°, 30°, 45°, 90° and 180°) and experiment 5 (−90°, -45°, -30°, -15°, 0°, 15°, 30° and 180) to compare the influence of varying the mean direction (mean upwards biased in experiment 4 and mean downwards bias in experiment 5). To compare the influence of prior movement direction on the magnitude of the serial dependence effect, we again computed the difference between trials with a fast and slow prior for the oculomotor response in the probe trial for each of the different directions (see Figure 4B for some example conditions from experiment 4).

**Figure 3.**
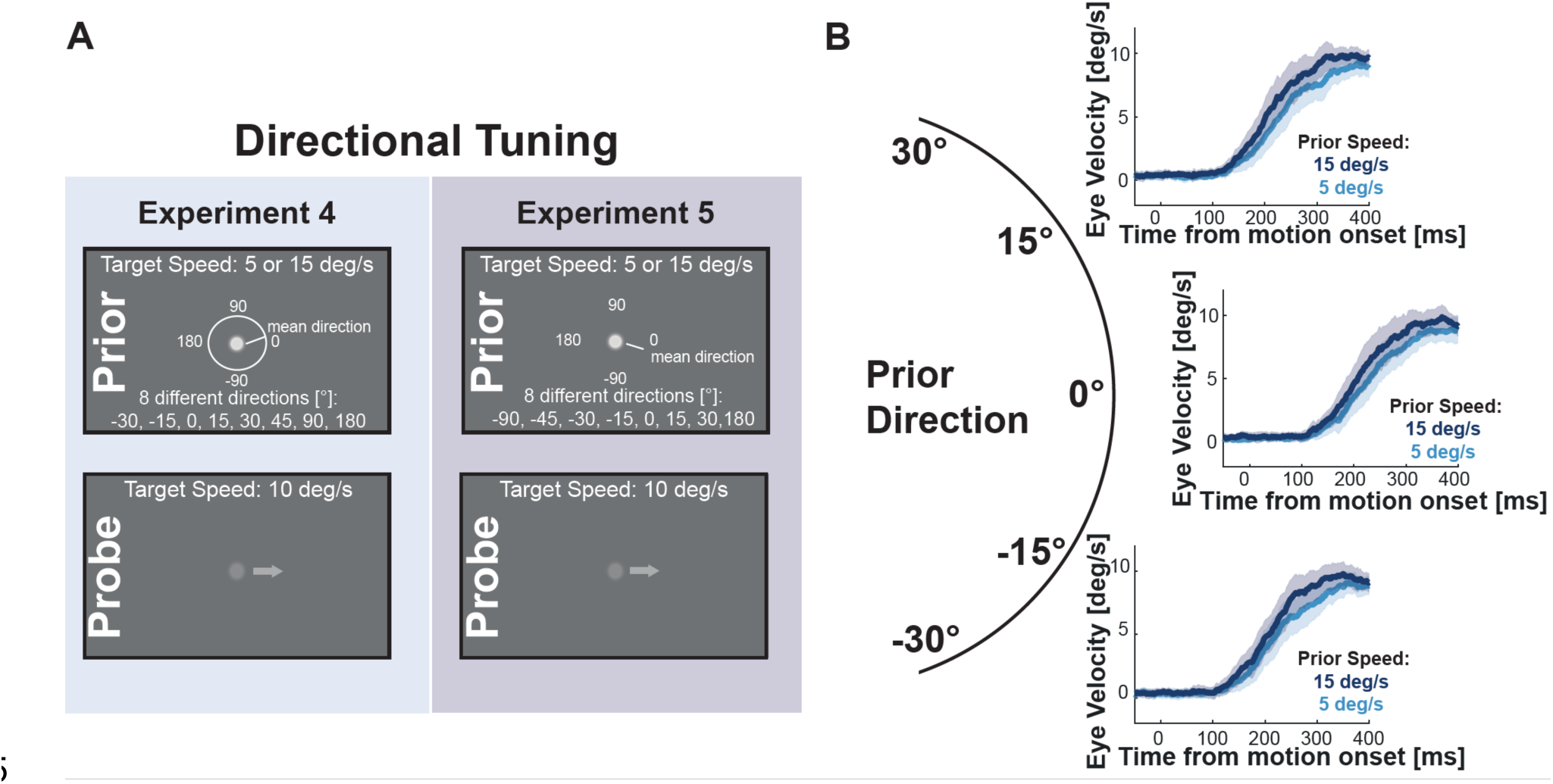
Directional tuning of serial dependence. **A** Depiction of the paradigms in the directional tuning experiments. The prior and probe stimulus always started at the same spatial location, but the relative movement direction between them differed. While the probe always moved horizontally to the right, the prior could have one of eight different directions. The mean direction of the prior movement differed between experiment 4 and 5: while the mean direction in experiment 4 was biased slightly upwards, the mean direction in experiment 5 was biased slightly downwards. **B** shows the average eye velocities across observers for the probe movement for experiment 4. Depicted is a selection of three different prior direction for -15°, 0° and 15° and the different shades of blue indicate the velocity of the prior movement. Please note that a prior direction of 0°, is identical to the collinear direction in the spatial tuning experiments. The shaded area depicts the 95% CI across observers.

**Figure 4.**
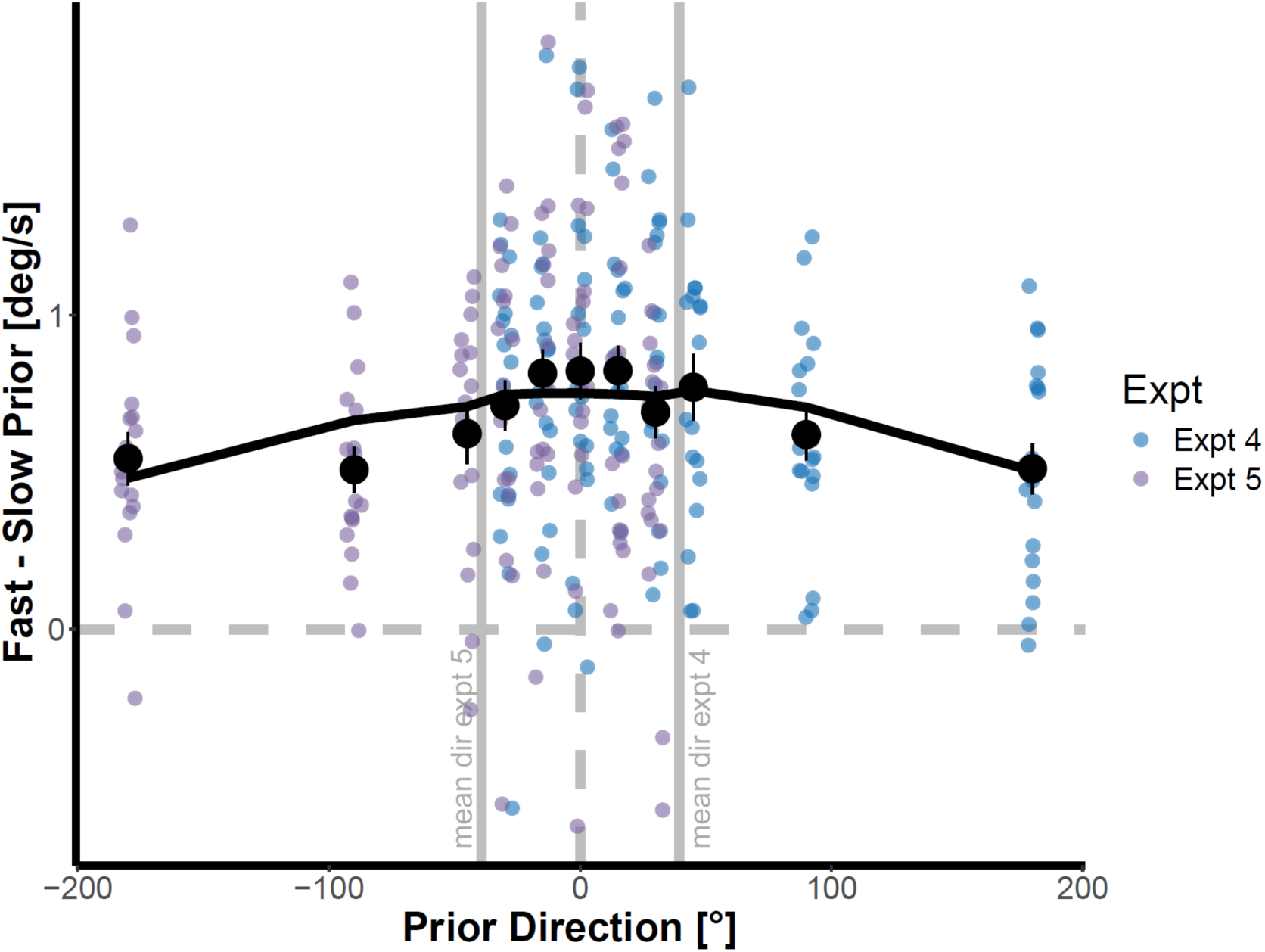
The magnitude of serial dependence depends on the direction of the prior. Small points represent individual participants for experiment 4 (blue) and experiment 5 (purple). Large black points represent mean serial dependence effect for each relative angle between prior and probe. The black line represents fitted growth curve model with a quadratic term.

### Methods – Directional tuning

#### Observer characteristics

The same group of 18 observers that completed Experiment 1, also completed Experiment 4. Another group of 18 observers (mean age: 21.54; range: 19-24; 14 female) took part in Experiment 5. All participants reported normal or corrected-to-normal vision and were naïve with respect to the study. Experimental procedures were in line with the declaration of Helsinki and were approved by the local ethics committee. Written informed consent was obtained from each participant.

#### Setup

The setup was the same as the spatial tuning experiments.

#### Experimental conditions

Experiments 4 & 5 were designed to specifically test the directional tuning of the serial dependence effect. The paradigm was the same as in the spatial tuning experiments: in each trial observers again saw two consecutive movements, however this time we systematically varied the direction of the prior movement with respect to the probe. The probe movement was identical across all trials and always started in the center and moved at 10 deg/s to the right (0°). The prior movement also started from the center, but could move in 8 different directions and again with two different speeds (5 deg/s or 15 deg/s). The movement directions were -30°, -15°, 0°, 15°, 30°, 45°, 90° and 180 in experiment 2, and -90°, -45°, -30°, -15°, 0°, 15°, 30° and 180 in experiment 3. 0° reflects a movement to the right, 90° a movement upwards, +/-180° a movement to the left and -90° a movement downwards. This allowed us to investigate how the different directions influenced the impact of the prior speed on tracking behavior in the probe trial. The timing of the presentation of the different movements was identical to experiment 1. Observers again completed 128 trials (8 prior directions * 2 prior speeds * 8 repetitions) and a total of 3 blocks, each lasting between 15 and 20 minutes. By comparing the serial dependence effect across the different relative directions, we could assess whether the serial dependence effect gets stronger the more the directions matched.

Since the same group of observers completed Experiment 1 and 4, the order of Experiment 1 and 4 was counterbalanced. Experiment 5 was collected in a different group of observers after Experiment 4 to assess the influence of the average direction in a given experiment.

#### Analysis

Preprocessing and identification of the serial dependence effect was the same as for the spatial tuning experiments. In addition, since in Experiments 4 & 5 we systematically varied the relative direction of prior and probe, we analyzed the initial direction of the pursuit response. Similar to our velocity measurement, we computed the angle of the eye between the eye position at motion onset and the eye position 400 ms after motion onset.

We again computed the serial dependence effect for eye velocity by looking at the difference in the horizontal eye velocity for the probe movement depending on the prior velocity (see also spatial tuning experiments). We computed this measurement for each of the 8 different prior orientations presented in each experiment. To look at the effect on the initial eye direction, we looked at the average angle computed for the probe movement in dependence of the prior direction. To account for individual biases in the initial eye angle, we subtracted the average angle for the 0° and 180° condition from all other directions, since in these conditions the prior should not have an influence, given that the prior and probe move along the same horizontal axis. To be able to compare the results across prior conditions and experiments, we then expressed the eye angle as proportion of adjustment from the relative difference (e.g. an angle of 3° for a prior moving target at 30° would be 0.1).

In an additional step we tried to relate the strength of the serial dependence effect on eye velocity with the serial dependence effect on eye direction across observers. We first estimated the reliability of the individual differences for these two measures. *For eye velocity,* for each subject we split all available trials for a respective condition into two random groups, 1000 times, and then for each of these iterations computed the serial dependence effects by the difference in the probe eye velocity between the fast and slow prior for each prior orientation. To isolate the retinal effect, we then went on two subtract the mean serial dependence effect across iterations for the 180° prior condition for the eye velocity. We then took the mean of the serial dependence effects across the remaining prior orientations (except 180°), and in this way obtained 1000 estimates for each of the two random groups for the velocity serial dependence effect for each observer.

*For eye direction*, we again split the trials per prior orientation into two random groups and for each of 1000 iterations computed the average eye angle during the probe, for each of the prior orientations for the two different groups. To correct again for the baseline differences in eye angle, for each of the iterations we subtracted the mean of the average eye angle for 0° and 180°, which then gave us separate estimates for each of the 1000 iterations for the directional serial dependence effect for each observer for each prior direction. We then averaged across the directions by again turning the eye angles into the proportion of the presented prior angle (see above) and averaged across the orientations. This again left us with 1000 estimates for each of the two random groups for the directional serial dependence effect for each observer. To estimate the split-half reliability, we collapsed all observers across both experiments and computed the correlation between the estimates obtained for the first random group and the second random group.

To now test the relationship between the velocity and directional serial dependence effect, we correlated the same metrics as used for the reliability estimates, but based on the whole group of trials for each of the respective conditions for each observer.

#### Exclusion criteria

We used the same exclusion criteria as for the spatial tuning. Based on these criteria we could include 6794 out of 6912 trials (98%) for experiment 4. For experiment 5, we had to exclude one observer due to a very high-exclusion rate (> 30%) and for the remaining 17 observers 6215 out of 6528 trials (95%) were included in the analysis.

#### Statistical Analysis

Growth curve modeling was conducted in R using the lme4 package (Bates et al., 2015). Pearson correlations were conducted in Matlab 2021b.

## Results – Directional Tuning

### The modulation of eye velocity

To assess whether the relative movement direction influenced the serial dependence effect (see Figure 4), we again computed the serial dependence effect as the difference between the fast and slow prior for the probe velocity. We included data from both experiments in the following analyses, given that there were no differences between Experiments 4 and 5 in the strength of the serial dependence effect in the conditions that were tested in both experiments (one-way ANOVA (serial_dep ∼ expt): F(1,173) = 1.044, p = 0.308). We used growth curve modelling (Mirman, 2017; Stewart et al., 2019; Stewart & Schütz, 2018) to quantify the linear and quadratic relationship between relative movement direction and serial dependence effect. We defined a baseline model with a linear term only (serial_dep ∼ rel_mov_dir), and compared this to a model with both linear and quadratic terms. The quadratic term was a second-order orthogonal polynomial fitted over relative movement direction (serial_dep ∼ rel_mov_dir(linear) + rel_mov_dir(quadratic)). Both models included participant as a random effect (random intercept only). Model comparisons revealed that adding the quadratic term significantly improved model fit: X^2^ = 12.96, p = 0.00032, AIC_linear = 321.19, AIC_quadratic = 310.23. Within the quadratic model, there was no significant linear relationship between serial dependence and relative movement direction: F(1,277.49) = 0.099, p = 0.75; however there was a significant quadratic relationship: F(1,244.86) = 13.30, p = 0.00032 (see Figure 4 for model fit). This suggests that when the relative movement directions were closer to zero, the serial dependence effect was highest, and the magnitude of this effect decreased as the prior direction deviated further from zero. The direction-unspecific component observed for the spatial tuning was also observed in these directional tuning experiments: For both experiments, we observed a significant attractive serial dependence effect even when the prior and probe moved in opposite directions.

### The modulation of eye direction

In a second step of the analysis, we assessed whether prior target direction also influenced the direction of the current oculomotor response. To visualize this influence, we looked at the average trajectories for the probe oculomotor responses during the first 400 ms after target motion onset, based on the relative prior directions (see Figure 6 A & B). Since observers had individual biases in how they tracked target movements, we subtracted the average vertical position across time for the 0° condition to account for these. Just by looking at the trajectories, one can see that the vertical position of the eye was affected by the relative direction of the prior (see Figure 5A and 5B). Prior targets moving downwards also led to a more downwards response in the probe trial and vice versa. We quantified this effect by computing the initial angle of the oculomotor response in this given interval (see Methods for more details). We again normalized the angle by subtracting the average angle for the 0° and 180° prior for each subject (See Figure 5C and 5D).

**Figure 5.**
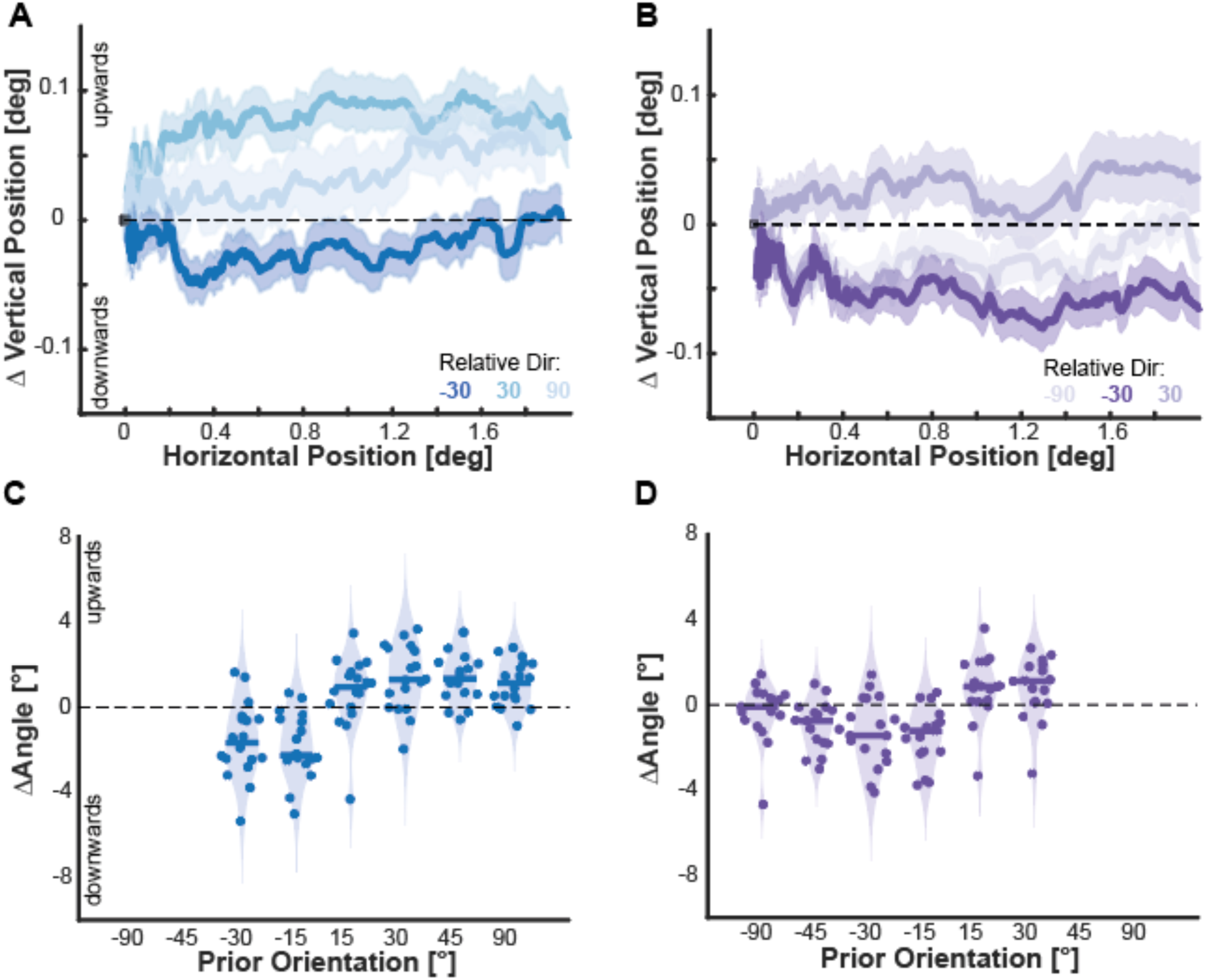
The influence of prior direction on eye direction. **A** and **B** show the average eye position trace in the first 400 ms after target movement onset for experiment 4 (A) and 5 (B). The shaded area depicts the standard error across observers. The different shaded of blue (A) or purple (B) depict a selection of the different relative directions of the prior target movement. **C** and **D** show the difference in the computed angle of the eye movement for all different prior directions and the prior movement moving to 0°. C shows the results for experiment 4, D shows the results for experiment 5. Negative values indicate a downward movement, positive values an upward movement. For each prior direction, the figure shows the distribution across observers with a violin plot, the solid line depicts the median and the dots depict the individual values per observer.

**Figure 6.**
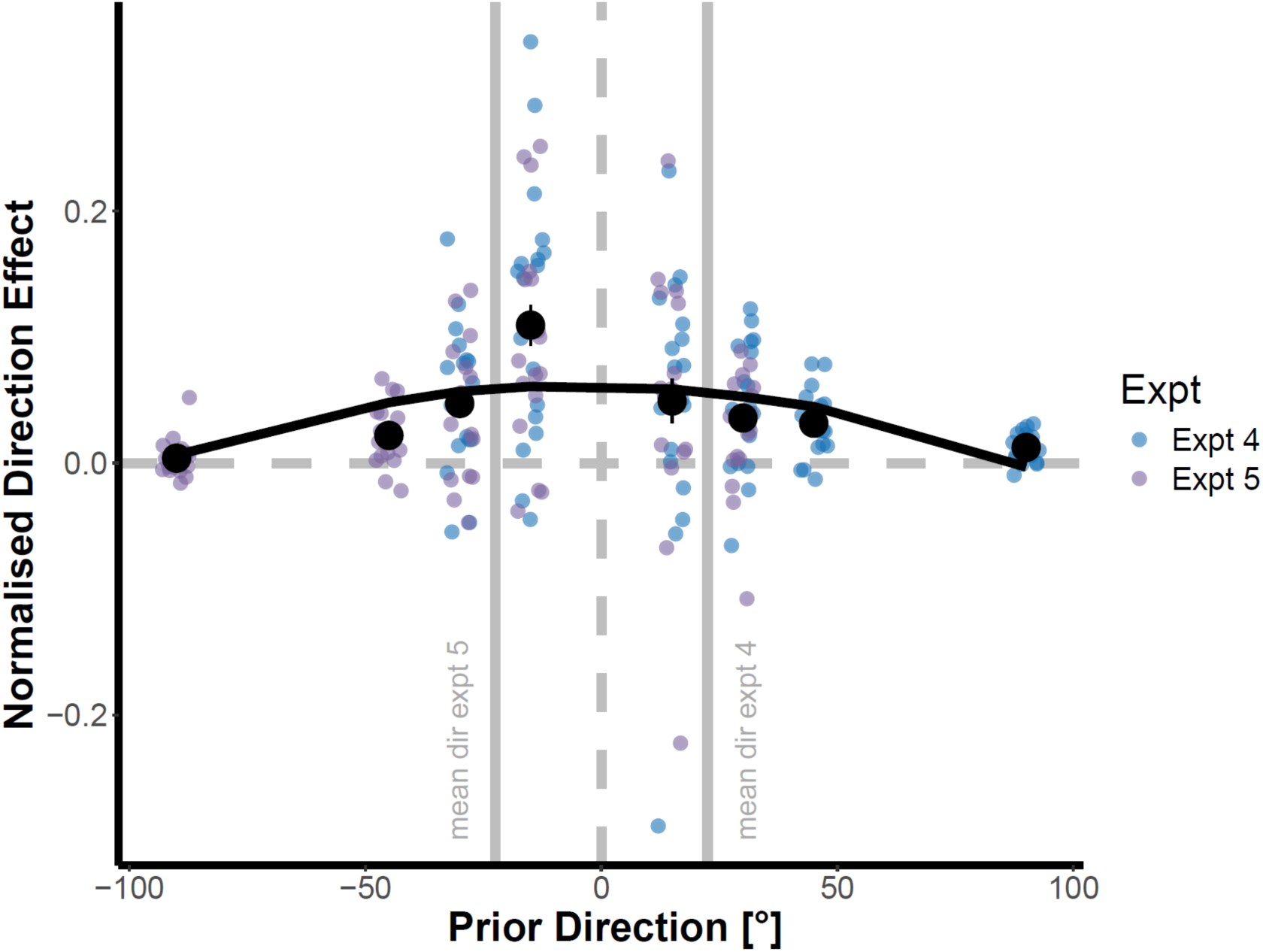
The influence of prior direction on the serial dependence effect. Small points represent individual participants for experiment 4 (blue) and experiment 5 (purple). Large black points represent mean serial dependence effect for each relative angle between prior and probe. The black line represents fitted growth curve model with quadratic term.

Next, we wanted to determine whether the direction of the prior affected the strength of the serial dependence effect. To compare the effect across the different prior angles, we normalized the direction effect by the magnitude of the prior angle. In this way, our measurement expresses the proportion of the angle difference that is transferred to the next trial (e.g. a 7.5 deg change in initial eye angle for a prior of 15 deg leads to a value of 0.5). We again collapsed across experiments (Kruskal-Wallis test (serial_dep ∼ expt): X^2^ = 1.043, p = 0.307) and used growth curve modelling to quantify the linear and quadratic relationship between prior direction and serial dependence effect. We defined a baseline model with a linear term only (serial_dep ∼ prior_dir), and compared this to a model with both linear and quadratic terms. The quadratic term was a second-order orthogonal polynomial fitted over prior direction (serial_dep ∼ prior_dir(linear) + prior_dir(quadratic)). Both models included participant as a random effect (random intercept only). Model comparisons revealed that adding the quadratic term significantly improved model fit: X^2^ = 17.72, p < 0.0001, AIC_linear = -496.88, AIC_quadratic = -512.6. Within the quadratic model, there was no significant linear relationship between serial dependence and prior direction: F(1,170.39) = 0.34, p = 0.56; however there was a significant quadratic relationship: F(1,98.78) = 17.42, p < 0.0001 (see Figure 6 for model fit). This suggests that when the prior horizontal movement direction was closer to zero, the serial dependence effect was highest, and the magnitude of this effect decreased as the relative angle between probe and prior increased. Together, these results show that not only the previous target speed but also the previous target direction transfers to the next trial. The size of this effect is again modulated by the relative difference in orientation.

### The relationship between the directional and velocity effect

Since we observed a significant influence on eye speed as well as eye direction, in the last step of the analysis we wanted to test the relationship between these two serial dependence effects. For each observer, we computed the magnitude of the retinal tuning effect for speed (see Figure 4) as the average over all different prior orientations. For the magnitude of the directional tuning effect, for each observer, we used the directional tuning and averaged across the prior directions and speeds (see Figure 6). Before we computed the correlation between the metrics, we estimated the reliability of the individual differences via a split-half correlation and estimated the variance via Bootstrapping (Figure 7A, see Methods for more details). To estimate the relationship between the velocity and directional serial dependence effect, we computed a Pearson correlation between the two individual estimates across all observers from both experiments. We obtained good (Median Correlation = .53 for Direction) to very good (Median Correlation = .78 for Speed) reliability estimates. Between the two metrics, we observed a significant correlation between the strength of the serial dependence for speed and direction (r(33) = .41, p = .0151): observers with a stronger serial dependence effect for speed also showed a stronger serial dependence effect for direction (Figure 7B). Note here, that when we compute the correlation between the retinal direction effect with the overall effect on eye velocity (including the retinal and un-specific component), this correlation disappears (r(33) = .04, p = .814) Thus, the two retinal factors do not only interact, but also seem to share some similarities.

**Figure 7.**
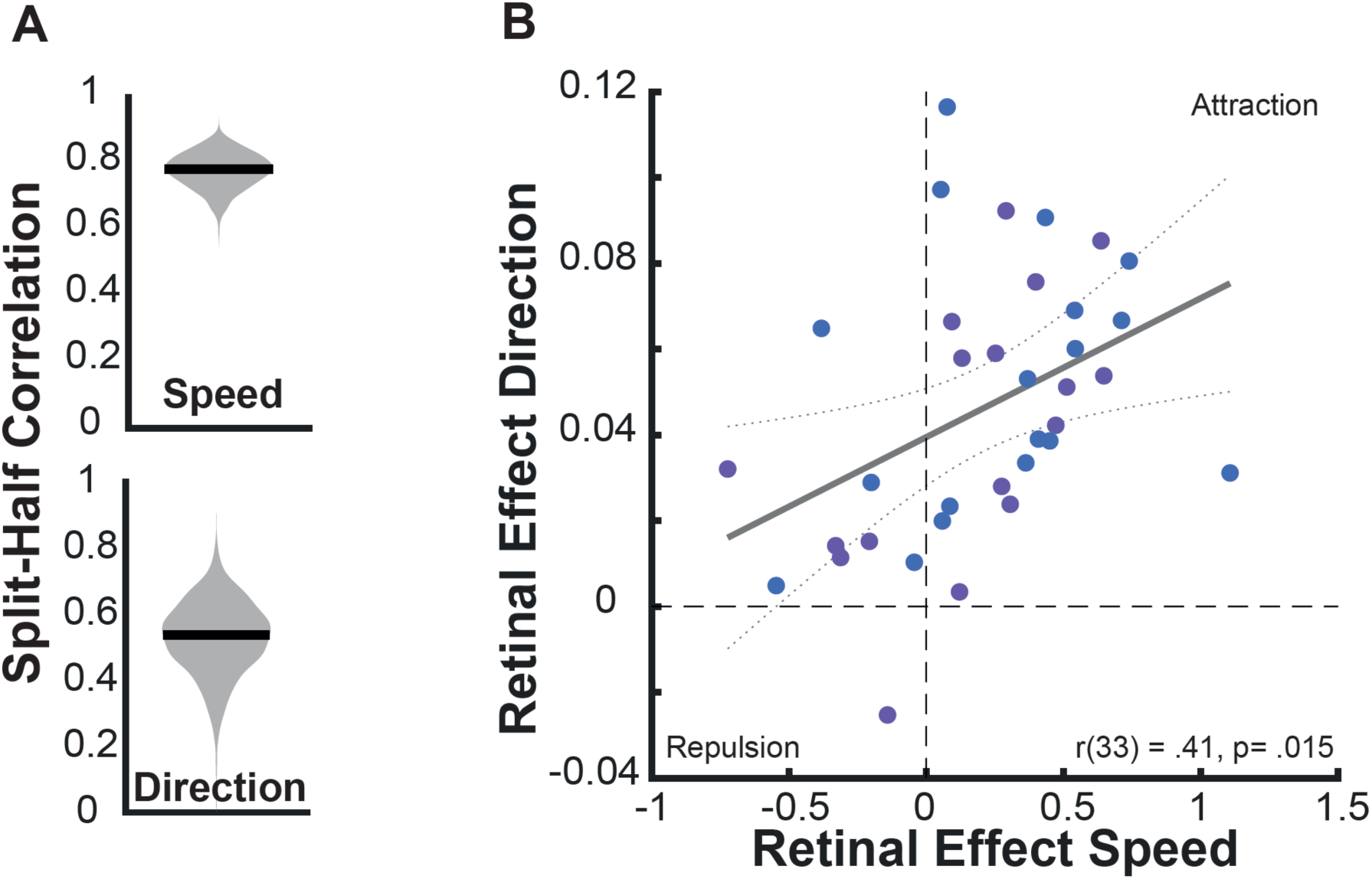
Relation between direction and velocity effect. **A** shows the estimated reliability of the individual differences we observed for the retinal target speed (upper panel) and target direction effect (lower panel). The estimate was done via Bootstrap (see Methods for more details) and shown is the distribution of obtained split-half correlation for 1000 different samples. The solid black line depicts the median split-half correlation. **B** The correlation between the strength of the retinal speed and retinal direction effect. Each data point depicts a single observer, data points in blue show observers from experiment 4 and data points in purple observers from experiment 5. The gray line depicts a linear regression through the data and the dashed line the 95% CI of this regression.

## Discussion

The goal of this study was to investigate the spatial and directional tuning of serial dependence for oculomotor tracking. In a first set of experiments, we observed that the serial dependence effect occurs mainly in a retinotopic reference frame, since there was no modulation of the strength of the effect, even when the prior and the probe stimulus were separated by 30 deg in spatiotopic coordinates (Figure 2). In line with this result, we found a modulation of the strength of the effect by target movement direction, with a stronger effect for collinear movements than for movements in the opposite direction. Importantly, for priors that moved into the opposite direction to the probe, we still observed a significant attractive serial dependence effect, which we attribute to a non-retinal component (Figure 2). In a second set of experiments, we investigated the directional tuning of serial dependence in more detail. Here we observed that the strength of the serial dependence on eye velocity (Figure 4) as well as eye direction was modulated by the relative angle between prior and probe (Figure 6). The effect was stronger for prior directions more similar to the probe direction. The strength of the effect on eye velocity and eye direction was correlated (Figure 7), with observers who showed a stronger influence of the prior on eye velocity also showing a stronger influence of the prior on eye direction.

### The spatial tuning of serial dependence for oculomotor control

For perceptual serial dependence, information is integrated if it falls within a certain spatial and temporal proximity to the current stimulus – the so-called serial dependence ‘continuity field’ (Fischer & Whitney, 2014). How this spatial proximity is computed is still an open question. Previous studies that measured the continuity field found that information separated by up to 20 deg can be integrated (Manassi et al., 2023), however in most of these studies the retinotopic and spatial location were confounded. Only a few studies have tried to dissociate retinotopic from spatiotopic serial dependence, and they observed ambiguous evidence. Mikellidou and colleagues (2021) dissociated the two reference frames by introducing different head positions and observed that mostly allocentric, spatiotopic cues could explain their results. In contrast, and in-line with the results of this study for oculomotor control, Collins (2019) observed that when the mismatch between the two reference frames occurs due to different fixation positions, then retinotopic information plays a larger role. One could speculate that head movements emphasize the need to use spatiotopic instead of retinotopic information (see Mikellidou et al. (2021) for a more detailed discussion of these discrepancies), but one important point made by Mikellidou and colleagues is that due to constantly changing retinal input due to eye and head movements, a perceptual serial dependence effect should be spatiotopic.

In contrast to serial dependence for perception, the critical information used for serial dependence by the oculomotor system seems to be early retinal signals (Goettker & Stewart, 2022): the goal of the oculomotor system is to bring and keep objects of interest on the fovea, and therefore the signals that should be integrated are retinal error signals like position or velocity error (Goettker & Gegenfurtner, 2021). Observing strong evidence for a retinotopic serial dependence effect for oculomotor tracking is consistent with the goals of the oculomotor system. The strength of the serial dependence effect was not affected by the distance between the prior and the probe stimulus, even for distances far larger than the 20 deg estimated to be the limit for perceptual integration. This retinotopic reference frame also matches the presumed neuronal underpinning of this effect (Darlington et al., 2018). The integration of previous experience seems to be based on an incoming velocity error signal from the middle temporal area (MT) which is then integrated with previous experience in the frontal eye field (FEF). While there is some evidence that MT can show spatiotopic characteristics (d’Avossa et al., 2007), there is also evidence for retinotopic coding (Gardner et al., 2008), especially if attending a moving target (Crespi et al., 2011).

### The directional tuning of serial dependence for oculomotor control

Our results replicate and extend the previous results from Darlington and colleagues (2017), who also observed that two characteristics of the prior target movement (direction and speed) both influence the following eye movement response. The shape of the relationship showing the influence of prior target direction on the magnitude of the serial dependency effect is consistent with a hypothesis that the signal mediating serial dependence for oculomotor control originates in area MT. We can speculate that the modulation of the serial dependence effect resembles the tuning curve of neurons with different preferred speeds or target movement directions (Albright, 1984; Born & Bradley, 2005; Dubner & Zeki, 1971).

Similar to Darlington and colleagues, we also observed that previous experience can not only change the current eye velocity and direction, but that these effects might be related to a common in a gain mechanism. Sensorimotor gain is the amplifier that defines how strongly a given sensory signal is driving the oculomotor system and this gain scales with eye velocity (Lee et al., 2013; Tanaka & Lisberger, 2002). For example, during pursuit the oculomotor system responds stronger to a visual perturbation than during fixation (Yang et al., 2012). In line with this idea, a higher gain for a faster prior target movement can also explain why despite more sensory noise, a faster prior target movement leads to a stronger directional effect for the subsequent movement (Darlington et al., 2017, 2018). Darlington et al explained their data by a model that is based on the modulation of a shared gain mechanism that is influencing the eye velocity and direction in the next trial. We observed additional behavioral evidence for this idea: there is a significant correlation between the retinal effects for eye direction and velocity across observers, which could reflect underlying differences in this shared gain mechanism (Figure 7). Interestingly, this relationship disappeared when we included the non-retinal component for the influence on eye speed, suggesting that this component might be independent of the gain control mechanism.

### A space and direction-independent serial dependence effect for oculomotor control

Previous studies have mostly tested serial dependence effects for oculomotor tracking with targets moving in the same or similar directions for the prior and the probe (Darlington et al., 2017; Deravet et al., 2018; Goettker, 2021). However, across all five experiments we observed one consistent and novel pattern: there was a significant positive serial dependence effect even when the prior and probe moved in opposite directions. This suggests that the serial dependence effect observed in most previous studies was probably a combination of at least two effects: a directionally-tuned retinal effect, and a more general, non-specific effect that is independent of the relative direction of the movement and which is not modified by spatial position. There could be two potential explanations for this effect: First, this could be due to some residual retinal motion signals, for example the motion arising from the background and screen border moving in the opposite direction of the screen target (Inaba et al., 2011). Second, the effect could reflect some cognitive component of the serial dependence effect potentially related to factors such as the level of attention (Hutton & Tegally, 2005) or working memory (Pascucci et al., 2023). These factors could work on a more general level that is independent of the direction of the prior stimulus, for example by leading to a more general difference in preparedness after faster stimuli, resulting in higher pursuit velocities.

While the retinal directionally tuned component seems to play a bigger role (see Figure 2), the non-specific component on average still accounts for roughly 40% of the effect, playing a substantial role in the overall magnitude of serial dependence. That serial dependence measurements can actually contain multiple components was also recently observed for perceptual judgements (Zhang & Alais, 2020). Zhang and Alais observed that individual differences in the strength of serial dependence effects were based on a different weighting of a positive choice and a repulsive motor bias.

## Conclusion

This study showed that serial dependence mainly occurs in a retinotopic reference frame, as the strength of the serial dependence was unaffected by a separation up to 30 deg in spatiotopic coordinates. Serial dependence was stronger for collinear movements compared to opposite directions, however even with movements in the opposite direction, a significant attractive serial dependence effect remained, indicating the existence of an additional non-retinal component. A more detailed investigation of directional tuning revealed that the strength of the serial dependence effect on eye velocity and direction was systematically influenced by the relative angle between prior and probe, with stronger effects for more similar directions. The prior angle modulated the serial dependence effect on eye velocity and eye direction, and the strength of these two effects was correlated. Overall, this study provides evidence for serial dependence in oculomotor control occurring in retinal coordinates, and furthermore hints at the presence of a secondary, non-retinal component in serial dependence for tracking eye movements.

## Data and Code Availability

Once the manuscript is accepted, all raw eye tracking data will be made available via OSF (osf.io/vj94n). The analysis code will be shared via Github.

## Acknowledgments

The authors would like to thank Zeynep Demirkan, Julia Schwab and Nils Borgerding for her help in data collection. This project was supported by the Deutsche Forschungsgemeinschaft (DFG), through project numbers 222641018-SFB/TRR 135 Project A1 (A.G.) and 460533638 (E.E.M.S.).

## Author contributions

Conceptualization, A.G. & E.E.M.S.; methodology, A.G. & E.E.M.S.; data collection, A.G.; formal analysis, A.G. & E.E.M.S.; writing—original draft, A.G. & E.E.M.S.; writing—review & editing, A.G. & E.E.M.S.; visualization, A.G. & E.E.M.S.; funding acquisition, A.G. & E.E.M.S.

